# Colocalization of GWAS and eQTL Signals Detects Target Genes

**DOI:** 10.1101/065037

**Authors:** Farhad Hormozdiari, Martijn van de Bunt, Ayellet V. Segrè, Xiao Li, Jong Wha J Joo, Michael Bilow, Jae Hoon Sul, Sriram Sankararaman, Bogdan Pasaniuc, Eleazar Eskin

## Abstract

The vast majority of genome-wide association studies (GWAS) risk loci fall in non-coding regions of the genome. One possible hypothesis is that these GWAS risk loci alter the individual’s disease risk through their effect on gene expression in different tissues. In order to understand the mechanisms driving a GWAS risk locus, it is helpful to determine which gene is affected in specific tissue types. For example, the relevant gene and tissue may play a role in the disease mechanism if the same variant responsible for a GWAS locus also affects gene expression. Identifying whether or not the same variant is causal in both GWAS and eQTL studies is challenging due to the uncertainty induced by linkage disequilibrium (LD) and the fact that some loci harbor multiple causal variants. However, current methods that address this problem assume that each locus contains a single causal variant. In this paper, we present a new method, eCAVIAR, that is capable of accounting for LD while computing the quantity we refer to as the colocalization posterior probability (CLPP). The CLPP is the probability that the same variant is responsible for both the GWAS and eQTL signal. eCAVIAR has several key advantages. First, our method can account for more than one causal variant in any loci. Second, it can leverage summary statistics without accessing the individual genotype data. We use both simulated and real datasets to demonstrate the utility of our method. Utilizing publicly available eQTL data on 45 different tissues, we demonstrate that computing CLPP can prioritize likely relevant tissues and target genes for a set of Glucose and Insulin-related traits loci. eCAVIAR is available at http://genetics.cs.ucla.edu/caviar/

## 1 Introduction

Genome-wide association studies (GWAS) have successfully detected thousands of genetic variants associated with various traits and diseases [1–4]. The vast majority of genetic variants detected by GWAS fall in non-coding regions of the genome, and it is unclear how these non-coding variants affect traits and diseases [5]. One potential approach to identify the mechanism of these non-coding variants on diseases is through integrating expression quantitative trait loci (eQTL) studies and GWAS[5]. This approach is based on the concept that a GWAS variant, in some tissues, affects expression at a nearby gene, and that both the gene and tissue may play a role in disease mechanism [6, 7].

Unfortunately, integrating GWAS and eQTL studies is challenging for two reasons. First, the correlation structure of the genome or linkage disequilibrium (LD) [8] produces an inherent ambiguity in interpreting results of genetic studies. Second, some loci harbor more than one causal variant for any given disease. We know that marginal statistics of a variant can be affected by other variants in LD [8–11]. For example, the marginal statistics of two variants in LD can capture a fraction of the effect of each other. Although GWAS have benefited from LD in the human genome by tagging only a subset of common variants to capture a majority of common variants, a fine mapping process, which attempts to detect true causal variants that are responsible for association signal at the locus, becomes more challenging. Colocalization determines whether a single variant is responsible for both GWAS and eQTL signals in a locus. Thus, colocalization requires correctly identifying the causal variant in both studies.

Recently, researchers proposed a series of methods [6, 12–17] to integrate GWAS and eQTL studies. PrediXscan [7] and TWAS [17] are examples of such methods, which impute gene expression followed by association of the imputed expression with trait. However, these methods do not provide a basis for determining colocalization of GWAS causal variants and eQTL causal variants. Another class of methods integrates GWAS and eQTL studies to provide insight about the colocalization of causal variants. For example, regulatory trait concordance (RTC) [13] detects variants that are causal in both studies while accounting for the LD. RTC is based on the assumption that removing the effect of causal variants from eQTL studies reduces or eliminates any significant association signal at that locus. Thus, when the GWAS causal variant is colocalized with the eQTL causal variant, re-computing the marginal statistics for the eQTL variant conditional on the GWAS causal variant will remove any significant association signal observed in the locus. Sherlock [12], another method, is based on a Bayesian statistical framework that matches the association signal of GWAS with those of eQTL for a specific gene in order to detect if the same variant is causal in both studies. Similar to RTC, Sherlock accounts for the uncertainty of LD. QTLMatch [16] is another proposed method to detect cases where the most significant GWAS and eQTL variants are colocalized due to causal relationship or coincidence. COLOC [14, 15], a method expanded from QTLMatch, is the state of the art method that colocalizes GWAS and eQTL signals. COLOC utilizes approximate Bayes factor to estimate the posterior probabilities for a variant that is causal in both GWAS and eQTL studies. Unfortunately, most existing methods for colocalization that utilize summary statistics assume presence of only one causal variant in any given locus for both GWAS and eQTL studies. As we show below, this assumption reduces the accuracy of results when the locus contains multiple causal variants.

In this paper, we present a novel probabilistic model for integrating GWAS and eQTL data. For each study, we use only the reported summary statistics and simultaneously perform statistical fine mapping to optimize integration. Our approach, eCAVIAR (eQTL and GWAS CAusal Variants Identification in Associated Regions), extends the CAVIAR [18] framework to explicitly estimate the posterior probability of the same variant being causal in both GWAS and eQTL studies while accounting for the uncertainty of LD. We apply eCAVIAR to colocalize variants that pass the genome-wide significance threshold in GWAS. For any given peak variant identified in GWAS, eCAVIAR considers a collection of variants around that peak variant as one single locus. This collection includes the peak variant itself, M variants that are upstream of this peak variant, and M variants that are downstream of this peak variant (e.g., M can be set to 50). Then, for all the variants in a locus, we consider their marginal statistics obtained from the eQTL study in all tissues and all genes. We only consider genes and tissues in which at least one of the genes is an eGene [19, 20]. eGenes are genes that have at least one significant variant (corrected p-value for multiple hypothesis of at least 10^−5^) associated with the gene expression of that gene. We assume that the posterior probability of the same variant being causal in both GWAS and eQTL studies are independent. Thus, this posterior probability is equal to the product of posterior probabilities for a given variant that is causal in GWAS and eQTL. We refer to the amount of support for a variant responsible for the associated signals in both studies as the quantity of colocalization posterior probability (CLPP).

Our framework allows for multiple variants to be causal in a single locus, a phenomenon that is widespread in eQTL data and referred to as allelic heterogeneity. Our approach can accurately quantify the amount of support for a variant responsible for the associated signals in both studies and identify scenarios where there is support for an eQTL mediated mechanism. Moreover, we can identify scenarios where the variants underlying both studies are clearly different. Utilizing simulated datasets, we show that eCAVIAR has high accuracy in detecting target genes and relevant tissues. Furthermore, we observe that the amount of CLPP depends on the complexity of the LD.

We apply our method to colocalize the Meta-Analyses of Glucose and Insulin-related traits Consortium (MAGIC) [21–24] GWAS dataset and publicly available eQTL data on 45 different tissues. We obtain 44 tissues from the GTEx eQTL dataset (Release v6, dbGaP Accession phs000424.v6.p1 available at: http://www.gtexportal.org) [19] and one tissue from the van de Bunt et al. (2015) study [25] that consists of human pancreatic islets tissue. Our results provide insight into disease mechanisms by identifying specific GWAS loci that share a causal variant with eQTL studies in a tissue. In addition, we identify several loci where GWAS and eQTL causal variants appear to be different. eCAVIAR is available at http://genetics.cs.ucla.edu/caviar/index.html

## 2 Results

### 2.1 Overview of eCAVIAR

The goal of our method is to identify target genes and the most relevant tissues for a given GWAS risk locus while accounting for the uncertainty of LD. Target genes are genes that their expression levels may affect the phenotype (e.g., disease status) of interest. Our method detects the target gene and the most relevant tissue by utilizing our proposed quantity of colocalization posterior probability (CLPP). eCAVIAR estimates CLPP, which is the probability that the same variant is causal in both eQTL and GWAS studies. eCAVIAR computes CLPP by utilizing the marginal statistics (e.g., z-score) obtained from GWAS and eQTL analyses, as well as the LD structure of genetic variants in each locus. LD can be computed from genotype data or approximated from existing datasets such as the 1000 Genomes data [26, 27] or HapMap [28]. We show in the Methods section that the marginal statistics of both GWAS and eQTL follow a multivariate normal distribution (MVN) given the causal variants and effect sizes for both studies. We use the MVN to estimate the CLPP. We show that CLPP is equal to the product of the posterior probability of the variant that is causal in GWAS and the posterior probability of the variant that is causal in eQTL. Computing the posterior probability of a causal variant is computationally intractable. Therefore, we assume a presence of at most six causal variants in a locus.

The estimated CLPP for a GWAS risk locus and a gene, which is obtained from eQTL studies, can be used to infer specific disease mechanisms. First, we identify genes that have expression levels affected by a GWAS variant. These genes are referred to as target genes. Second, we identify in which tissues the eQTL variant has an effect. To identify target genes, we compute CLPP for all genes in the GWAS risk locus. Genes that have a significantly higher CLPP in comparison are selected as target genes (Figure 1a). Similarly, we compute CLPP for all tissues and identify relevant tissues as those with comparatively high values of CLPP (Figure 1b). Figure 1b shows that the GWAS risk locus affects the Gene4 and the relevant tissues are liver and blood. However, Figure 1b indicates that pancreas is not a relevant tissue for this GWAS risk locus. Another application of CLPP is to identify loci where the causal variants between GWAS and eQTL studies are different. We can identify these loci if CLPP is low for all variants in the loci, and if there are statistically significant variants in both GWAS and eQTL studies.

**Figure 1.**
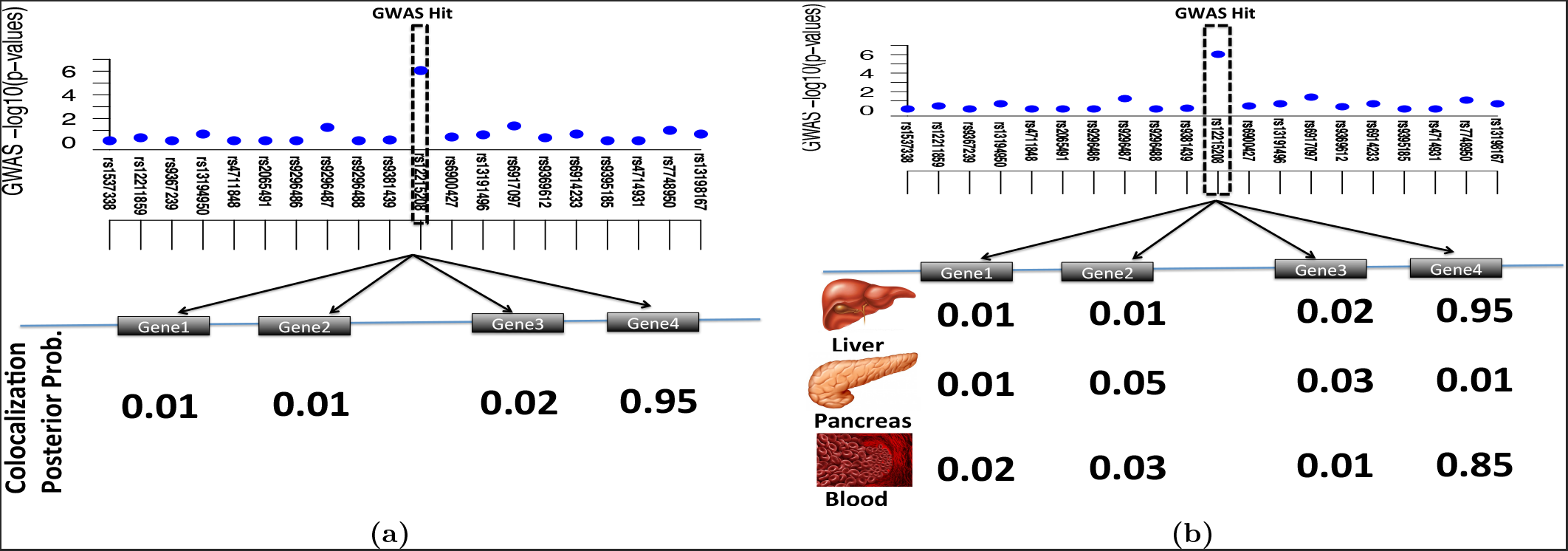
Overview of our method for detecting the target gene and most relevant tissue. We compute the CLPP for all genes and all tissues. Panel (a) illustrates a simple case where we only have one tissue and we want to find the target gene. We consider all the genes for this GWAS risk locus, and we observe Gene4 has the highest CLPP. Thus, in panel (a) the target gene is Gene4. In Panel (b), we have 3 tissues and we utilize the quantity of CLPP. Thus, the target gene is Gene4 again. Moreover, in this example, we consider that liver and blood are relevant tissues for this GWAS risk locus, while pancreas is not relevant to this GWAS risk locus.

To better motivate the behavior of CLPP, we consider the following four scenarios in Figure 2. In the first scenario, the same variant has effects in both GWAS and eQTL studies. Thus, its CLPP is high (Figure 2a). In the second scenario, we consider that the variant is associated with a phenotype in GWAS and not associated with gene expression. In this case, the quantity of CLPP is low (Figure 2b). In the third scenario, we consider that the variant is not associated with a phenotype in GWAS. However, it is associated with expression of a gene. In this case, CLPP is not computed for this variant. Rather, we compute CLPP for GWAS risk loci that are considered significant. In the fourth scenario, we have a variant that appears significant in both GWAS and eQTL. However, other variants in GWAS or eQTL are also significant due to high LD with the causal variant. The complex LD (see Figure 2c) of these variants results in a low CLPP. Here, we remain uncertain about which variants are actual causal variants. Finally, Figure 2d illustrates an example in which there is more than one causal variant. This demonstrates that underestimation of CLPP can result from assuming presence of a single causal variant. In this example, we have a locus with 35 variants (SNPs) and we have two causal variants (SNP6 and SNP26) that are not in high LD with each other. If we assume we have only one causal variant there are 35 possible causal variants for this locus and most of the causal variants have very low likelihood. The likelihood of two variants in which the SNP6 or SNP26 are selected as causal have similar likelihood and their likelihood is much higher than other variants. In this example, the estimated posterior probability of SNP6 or SNP26 being causal is equal to 50%. Thus, the estimated CLPP for SNP6 or SNP26 is 25%. However, if we allow more than one causal variant in the locus, all sets of causal variants have very low likelihood values except the set with both SNP6 and SNP26 selected as causal. In this case, the posterior probability of SNP6 or SNP26 being causal is close to 1. Thus, in this case we assume that we have more than one causal variant in this locus, the CLPP of SNP6 and SNP26 are close to 1.

**Figure 2.**
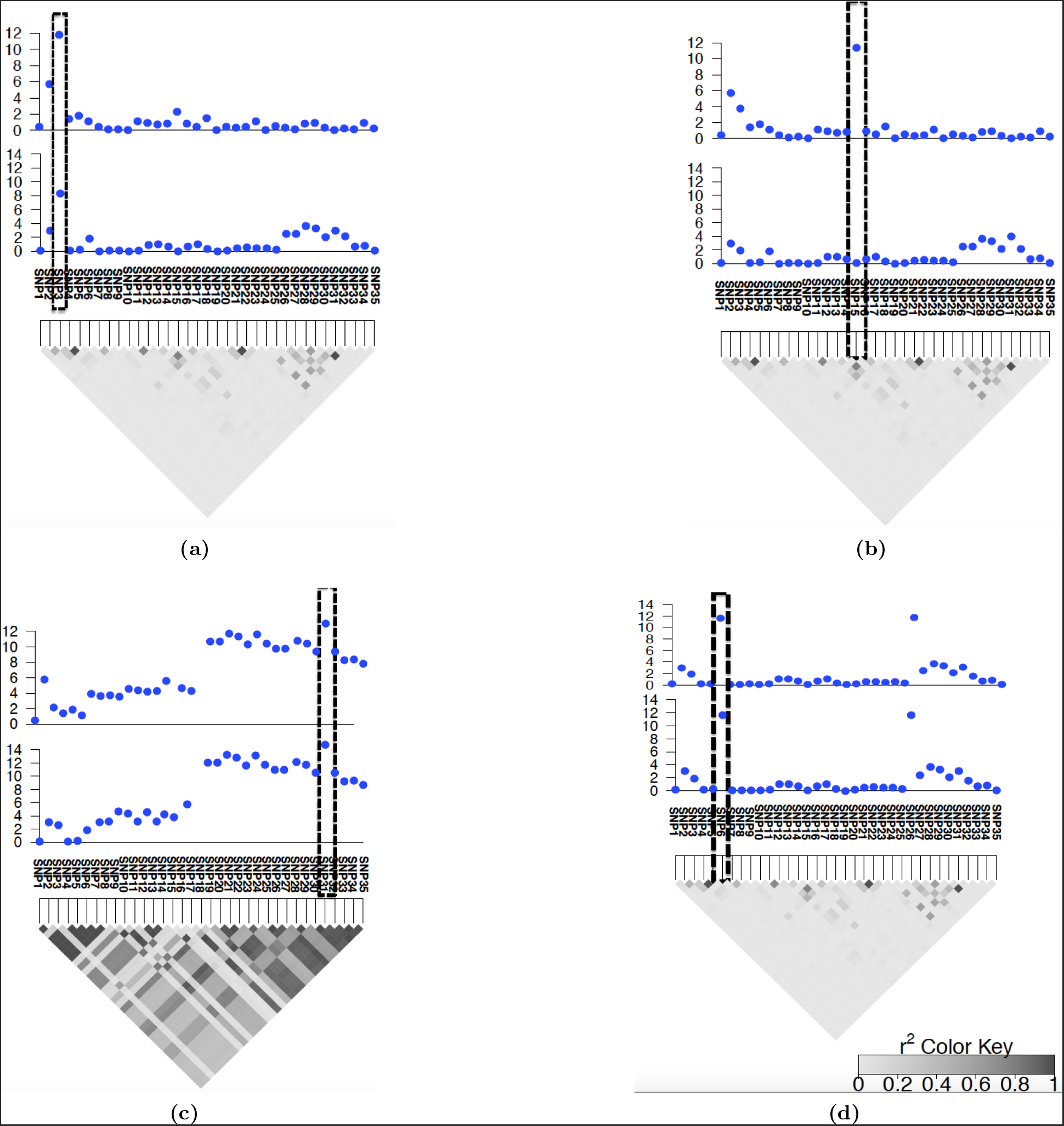
Overview of eCAVIAR. At high level eCAVIAR aligns the causal variants in eQTL and GWAS. The x-axis is the variant (SNP) location. The y-axis is the significant score (-log of p-value) for each variant. The grey triangle indicates the LD structure where every diamond in this triangle indicates the Pearson’s correlation. The darker the diamond the higher the correlation and the lighter the diamond the lower the correlation between the variants. The case where the causal variants are aligned the colocalization posterior probability (CLPP) is high for the variant that is embedded in the dashed black rectangle as shown in panel (a). However, the case where the causal variants are not aligned (the causal variants are not the same variants) then the quantity of CLPP is low for the variant that is embedded in the dashed black rectangle as shown in panel (b). In the case, the LD is high, which implies the uncertainty is high due to LD, the CLPP value is low for the variant that is embedded in the dashed black rectangle as shown in panel (c). Panel (d) illustrates a case where in a locus we have two independent causal variants. If we consider that we only have one causal variant in a locus, then the CLPP of the causal variants are estimated to be 0.25. However, if we allow to have more than one causal variant in the locus, eCAVIAR estimates the CLPP to be 1.

### 2.2 eCAVIAR Accurately Computes the CLPP

In this section, we use simulated datasets in order to assess the accuracy of our method. We simulated summary statistics utilizing the multivariate normal distribution (MVN) that is utilized in previous studies [18, 29–32]. More details on simulated data are provided in Section 3.4.2. In one set of simulations, we fix the effect size of a genetic variant so that the statistical power for the causal variant is 50%. In another set, we fix the effect size so that the power is 80%. We consider two cases. In the first case, we only have one causal variant in both studies. In the second case, we have more than one causal variant in these studies. For both cases, we simulated two datasets. In the first dataset, we implanted a shared causal variant. We generated 1000 simulated studies, which we then use to compute the true positive rate (TP). In the second dataset, we implanted a different causal variant in eQTL and GWAS. We filter out cases where the most significant variant is different between the two studies. Similarly to the previous case, we generated 1000 simulated studies.

#### 2.2.1 eCAVIAR is Accurate in the Case of One Causal Variant

We apply eCAVIAR to the simulated datasets and compute the CLPP for each case. We use different cut-offs to determine whether or not a variant is shared between two studies. For each cut-off, we compute the false positive rate (FP) and true positive rate (TP). The baseline method detects the most significant variant in a study as the causal variant. Thus, in the baseline method, we have colocalization when the most significant variant in both eQTL and GWAS is the same variant. We refer to this method as the Shared Peak SNP (SPS) method. The results are shown in Figures 3a and 3d Moreover, we plot the same results in receiver operating characteristic (ROC) curve (Figure S1). We observe our method has higher TP and lower FP compared to SPS. However, eCAVIAR has low TP when the cut-off for CLPP is high. Furthermore, eCAVIAR has an extremely low FP. Our results imply that eCAVIAR has high confidence for selecting loci to be colocalized between the GWAS and eQTL. eCAVIAR is conservative in selecting a locus to be colocalized. Given the high cut-off of CLPP, eCAVIAR can miss some true colocalized loci. However, loci that are selected by eCAVIAR to be colocalized are likely to be predicted correctly.

**Figure 3.**
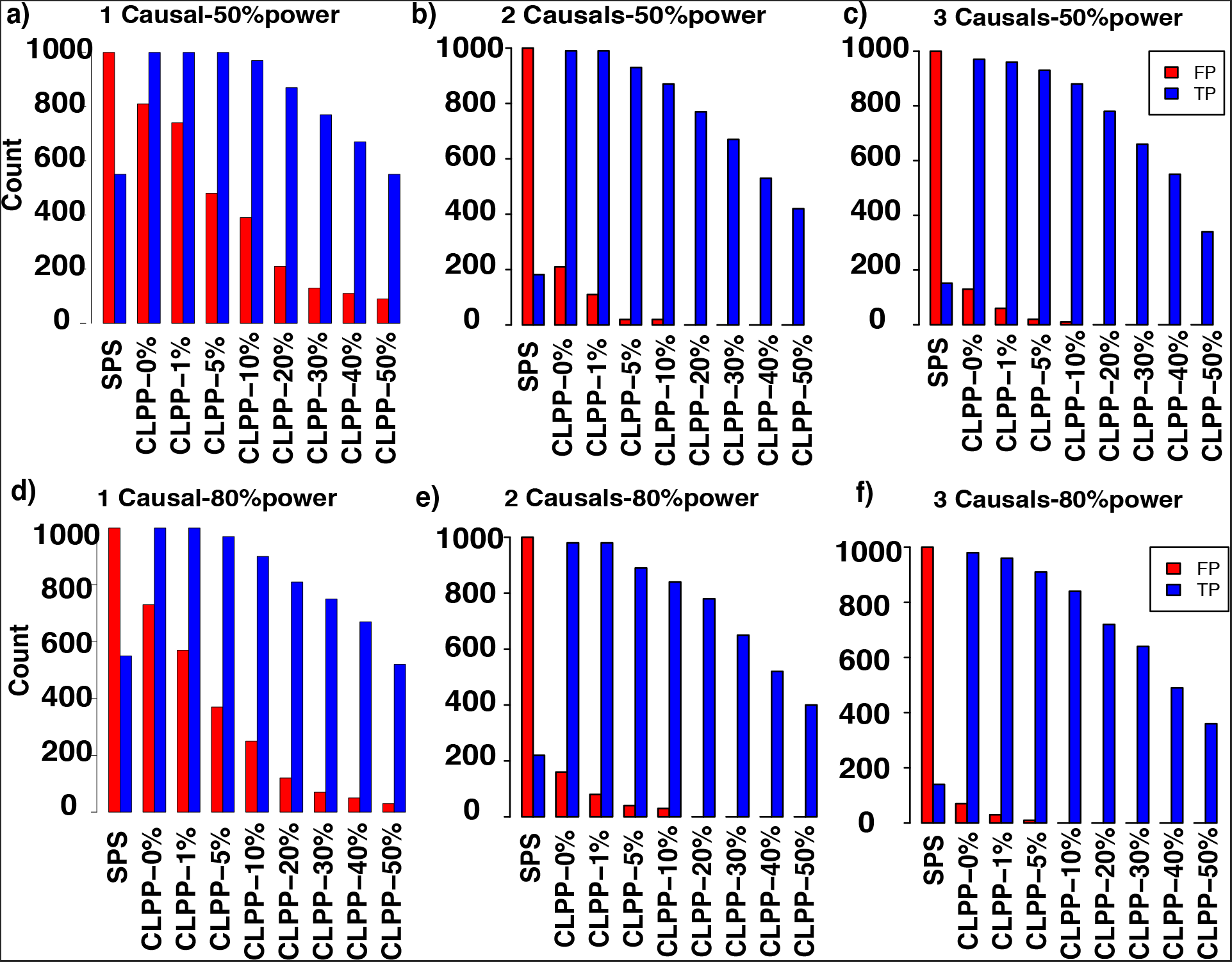
eCAVIAR is robust to the presence of allelic heterogeneity. We simulate marginal statistics directly from the LD structure for eQTL and GWAS. We implant one, two or three causal variants in both studies. Panels (a), (b), and (c) indicate the result for one, two, and three causal variants respectively where the statistical power on the causal variants is 50%. Panels (d), (e), and (f) indicate the result for one, two, and three causal variants respectively where the statistical power on the causal variants is 80%. eCAVIAR has a low TP for high cut-off, and eCAVIAR has low FP. This indicates that eCAVIAR has high confidence in detecting a locus to be colocalized between GWAS and eQTL, even in the presence of allelic heterogeneity.

The computed CLPP depends on the complexity of the LD at the locus. We apply eCAVIAR to the simulated datasets and compute the CLPP (Figure S2). Here, the average quantity of CLPP decreases as we increase the Pearson’s correlation (r) between paired variants. This effect increases complexity of LD between the two variants. Furthermore, the 95% confidence interval for the computed quantity increases as we increase the Pearson’s correlation. This result implies that the computed CLPP can be small for a locus with complex LD, even when a variant is colocalized in both GWAS and eQTL studies.

#### 2.2.2 eCAVIAR is Robust to the Presence of Allelic Heterogeneity

The presence of more than one causal variant in a locus is a phenomenon referred to as allelic heterogeneity (AH). AH may confound the association statistics in a locus, and colocalization for a locus harboring AH is challenging. In order to investigate the effect of AH, we perform the following simulations. We implanted two or three causal variants in both GWAS and eQTL, and we then generated the marginal statistics using MVN as mentioned in the previous section. Next, we compute TP and FP for eCAVIAR and SPS (see Figure 3). In the case of eCAVIAR, we consider a true colocalization when it can detect all the variants that colocalized. However, in the case of SPS, we consider a true colocalization when it can detect at least one of the variants that colocalized. Figures 3a, 3b, and 3c illustrate results of one, two, and three causal variants, respectively, when the statistical power is 50%. In a similar way, Figures 3d, 3e, and 3f illustrate results of one, two, and three causal variants, respectively, when the statistical power is 80%. Interestingly, SPS has a very low TP when there are two or three causal variants (see Figure 3). This implies that SPS is not accurate when AH is present. Similar to cases with one single casual variant (see Figures 3a and 3d), eCAVIAR has a very low FP when there are two or three causal variants (see Figures 3b, 3c, 3e, and 3f). This implies that eCAVIAR has high confidence in detecting a locus to be colocalized between GWAS and eQTL.

We generated simulated studies where the causal variants are different between the two studies. We computed the CLPP for all the variants in a region. Our experiment indicate that eCAVIAR has high TN and extremely low FN. eCAVIAR has high negative predictive value (NPV), NPV = *TN/(TN* + *FN*). These results are shown in Figures S3 and S4. Thus, eCAVIAR can detect with high accuracy loci where the causal variants are different between the two studies.

### 2.3 eCAVIAR is More Accurate than Existing Methods

We compare the results of eCAVIAR with RTC [13] and COLOC [14], two well known methods for eQTL and GWAS colocalization. We can use the previous section to generated simulated datasets; however, RTC is not designed to work with summary statistics. In order to provide a dataset compatible with RTC, we simulated eQTL and GWAS phenotypes under a linear additive model where we use simulated genotypes obtained from HAPGEN2 [33]. More details on the simulated datasets are provided in Section 3.4.3.

We compare the accuracy, precision, and recall rate of all three methods. Each method computes a probability for a variant to be causal in both eQTL and GWAS. In order to determine this probability for our comparison, we need to select two cut-off thresholds. We devised one threshold for detecting variants that are colocalized in both studies and another threshold to detect variants that are not colocalized. Here, we consider a variant to be causal in both studies if the probability of colocalization is greater than the colocalization cut-off threshold. The second cut-off threshold is used to detect variants that are not causal in both studies. We consider a variant is non-causal in both studies if the probability of colocalization is less than the non-colocalization cut-off threshold. In our experiment, we set the non-colocalization cut-off threshold at 0.1%, and for the colocalization cut-off threshold, we vary this value from 0.1% to 90%.

eCAVIAR outperform existing methods when the locus has one causal variant. We observe that all three methods have a similarly high recall rate (see Figure S5). eCAVIAR has much higher accuracy and precision in comparison to RTC (see Figure 4). Next, we consider the performance of the three methods when the locus has allelic heterogeneity. We use the same simulation described in this section, but in this case we implant two causal variants instead of one causal variant. In this setting, eCAVIAR has higher accuracy and precision when compared to COLOC and RTC. However, RTC has a slightly higher recall rate in comparison to eCAVIAR. Moreover, RTC tends to perform better than COLOC in the presence of allelic heterogeneity (see Figure 5). This result indicates eCAVIAR is more accurate than existing methods–even in the presence of allelic heterogeneity. However, if there exists only one causal variant in a locus, COLOC has better performance than RTC. In cases with more than one causal variant, RTC has better performance. These results are obtained when we set the non-colocalization cut-off threshold to be 0.1%. We change this value to 0.01% to check the robustness of eCAVIAR. We observe for different values of non-colocalization still eCAVIAR outperforms existing methods (see Figures S6 and S7). In all of the above experiments, we implant the causal variants uniformly in the locus. Next, we simulate causal variants in genomic variants that are enriched with functional annotations. In order to simulate the genomic enrichment, we use the same process that is utilized in PAINTOR [34]. We observe that eCAVIAR outperforms existing methods in these experiments (Figures S8 and S9).

**Figure 4.**
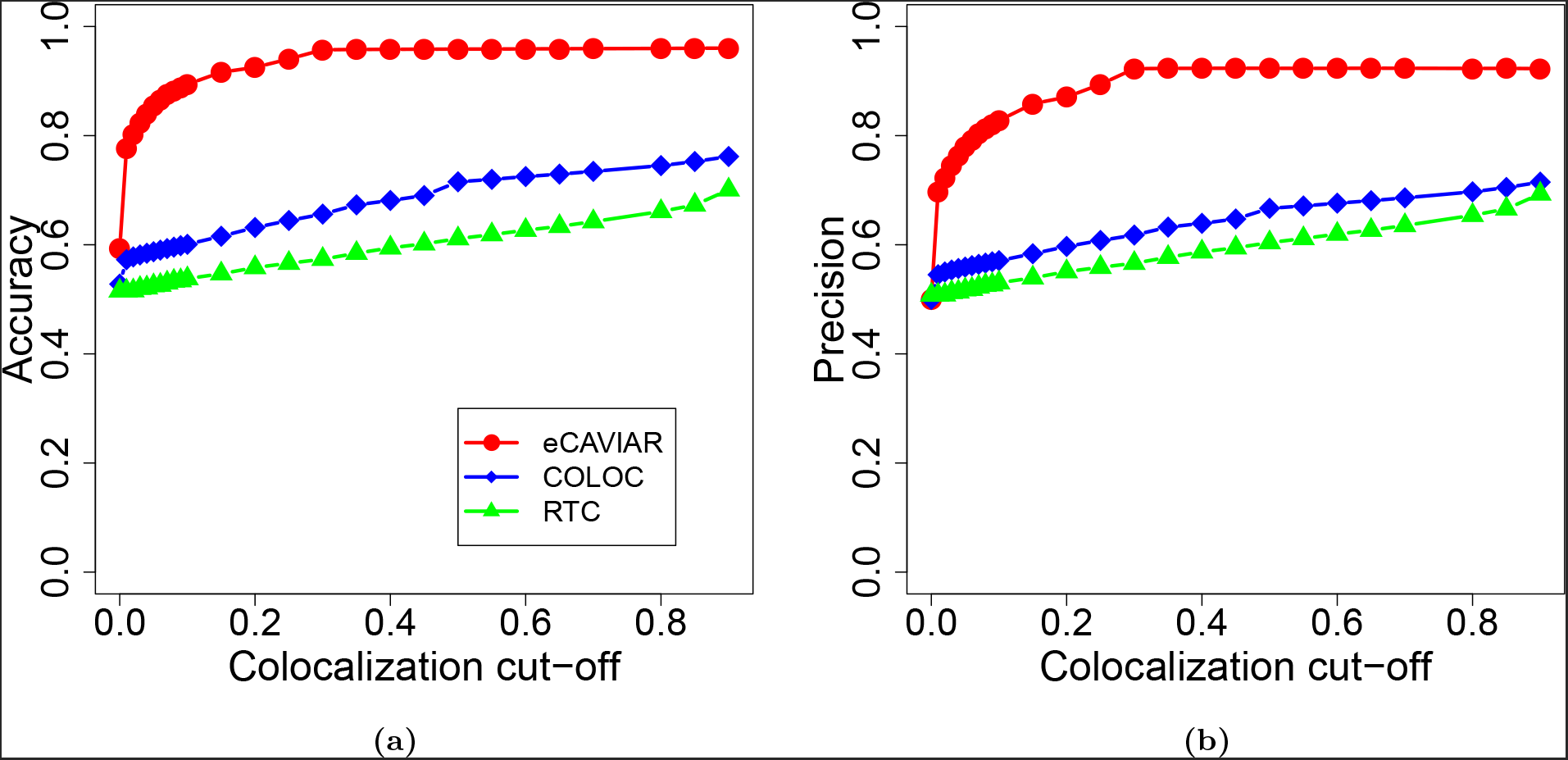
eCAVIAR is more accurate than existing methods for regions with one causal variant. We compare the accuracy and precision of eCAVIAR with the two existing methods (RTC and COLOC). The x-axis is the colocalization cut-off threshold. In these datasets we implant one causal variant, and we utilize simulated genotypes. We simulate the genotypes using HAPGEN2 [33] software. We use the European population from the 1000 Genomes data [26, 27] as the starting point to simulate the genotypes. Panels (a) and (b) illustrate the accuracy and precision respectively for all the three methods. We compute TP (true positive), TN (true negative), FN (false negative), and FP (false positive) for the set of simulated datasets where we generate the marginal statistics utilizing the linear model. Accuracy is the ratio of (TP+TN) and (TP+FP+FN+TN), 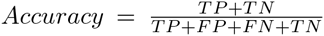, and precision is the ratio of TP and (TP+FP), 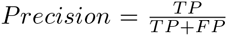. We set the non-colocalization cut-off threshold to 0.001. We observe eCAVIAR and COLOC have higher accuracy and precision compared to RTC.

**Figure 5.**
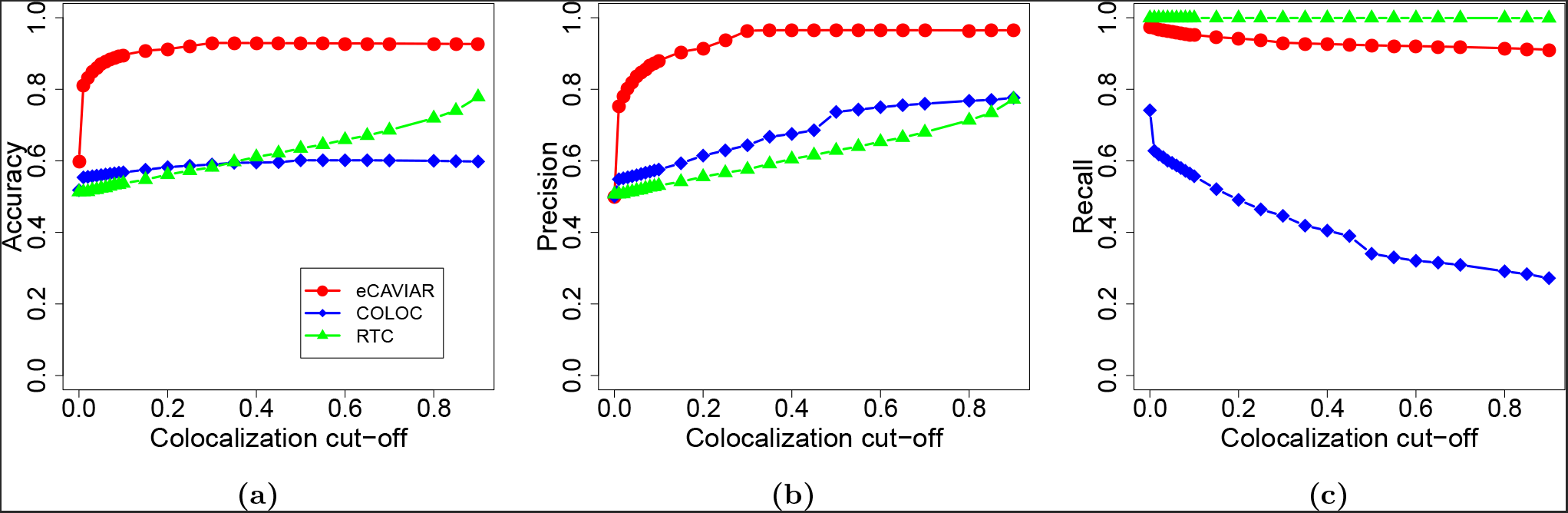
eCAVIAR is more accurate than existing methods in presence of allelic heterogeneity. We use similar process to generate the datasets as shown in Figure 4. However, in this case, we implant two causal variants. We simulate the genotypes using HAPGEN2 [33] software. We use the European population from the 1000 Genomes data [26, 27] as the starting point to simulate the genotypes. We compare the accuracy, precision, and recall rate. In these results, eCAVIAR tends to have higher accuracy and precision compared to the RTC and COLOC. However, RTC have slightly higher recall rate.

eCAVIAR has better performance when compared to COLOC and RTC, the pioneering methods for eQTL and GWAS colocalization. COLOC and RTC require different input data to perform the colocalization. COLOC only requires the marginal statistics from GWAS and eQTL studies. Unlike eCAVIAR, COLOC and RTC do not require the LD structure of genetic variants in a locus. However, RTC requires individual level data (genotypes and phenotypes) and is not applicable to datasets for which we have access to the summary statistics.

### 2.4 Effect of eQTL Sample Size on CLPP

We know that the statistical power to detect a true casual variant increases as we increase the number of samples in GWAS. As most GWAS sample sizes are in order of thousand of samples, we would like to investigate the effect of eQTL sample size on colocalization.

We simulate datasets where we set the number of samples in GWAS to be 5000. Then, we vary the number of samples in eQTL studies from 500 to 3500. We simulate the effect size for the causal variant in eQTL such that it accounts for the 0.01, 0.04, and 0.1 portion of heritability. We compute the CLPP for different cases; the distribution of CLPP are shown in box plot in Figure S10. The red horizontal line indicates the 0.01 (1%) colocalization cut-off used for eCAVIAR. We observe in the case where the causal variant accounts for 0.01 portion of heritability, we require at least 2000 samples for eQTL. Conversely, when the causal variant accounts for larger portion of heritability, eCAVIAR requires fewer samples.

### 2.5 Integrating Available eQTL for 45 Tissues and MAGIC Datasets Using eCAVIAR

We utilize the Meta-Analyses of Glucose and Insulin-related traits Consortium (MAGIC) dataset and GTEx dataset [19] to detect the target gene and most relevant tissue for each GWAS risk locus. MAGIC datasets consist of 8 phenotypes [21]. These phenotypes are as follows: FastingGlucose, FastingInsulin, FastingProinsulin, HOMA-B (*β*-cell function), HOMA-IR (insulin resistance), and Hb1Ac (Hemoglobin A1c test for Diabetes), 2-hour glucose, and 2-hour insulin after an oral glucose tolerance test. In our analysis, we use FastingGlucose (FG) and FastingProinsulin(FP) phenotypes containing the most number of significant loci. FG phenotypes have 15 variants and FP phenotypes have 10 variants that are reported significantly associated with these phenotypes from previous studies [21, 22]. We consider 44 tissues provided by the GTEx consortium (Release v6, dbGaP Accession phs000424.v6.p1 available at: http://www.gtexportal.org) [19], as well as previously published data on human pancreatic islets [25] - one of the key tissues in glucose metabolism, which is not captured in the GTEx data. Table S1 lists tissues and the number of individuals for each tissue.

We want to detect the most relevant tissue and a target gene for each of previously reported significant variants in GWAS. eCAVIAR utilizes the marginal statistics of all variants in a locus obtained from GWAS and eQTL. We obtain each locus by considering 50 variants upstream and downstream of the reported variant. Then, we consider genes where at least one of the variants in the locus is significantly associated with the gene expression of that gene. Thus, for one GWAS variant, there may exist multiple genes in one tissue that satisfy these requirements, and we consider these pairs of variants and genes as potential colocalization loci. Table S2 and Table S3 list the potential colocalization loci for FG and FP phenotypes, respectively. For any given variant, we use CLPP to detect the most relevant tissue and a target gene. We then select the gene and tissue that have highest CLPP as the target gene and the most relevant tissue, respectively.

Table 1 and Table 2 indicate the result of eCAVIAR for FastingGlucose and FastingProinsulin, respectively. This result shows genetic variants that are causal in both eQTL and GWAS. We only considered variants that are reported to be significant with FG [21] and FP [22] phenotypes. We use the cut-off threshold of 0.01 (1%) to conclude that two causal variants are shared.

**Table 1.** eCAVIAR joint analysis of FastingGlucose and GTEx dataset. We use *N* to indicate the number of individuals in each tissue that we have access to summary statistics from GTEx [19] and van de Bunt et al. (2015) study [25].

| chrom | pos. | rsID | Relevant Tissue | Target Gene |
| --- | --- | --- | --- | --- |
| 3 | 123082398 | rs11717195 | Islet ($N=118$) | *ADCY5* |
| 7 | 15064309 | rs2191349 | Islet ($N=118$) | *DGKB* |
| 7 | 44235668 | rs4607517 | Colon Sigmoid ($N=124$)<br>Thyroid ($N=278$) | *GCK*<br>*GCK* |
| 11 | 45873091 | rs11605924 | Whole Blood($N=338$) | *MAPK8IP1* |
| 11 | 47336320 | rs7944584 | Nerve Tibial ($N=256$)<br>Artery Tibial ($N=285$)<br>Islet ($N=118$)<br>Pituitary ($N=87$)<br>Artery Tibial ($N=285$)<br>Nerve Tibial ($N=256$)<br>Nerve Tibial ($N=256$) | *CELF1*<br>*MADD*<br>*MADD*<br>*MADD*<br>*MDK*<br>*NR1H3*<br>*RAPSN* |

**Table 2.** eCAVIAR joint analysis of FastingProinsulin and GTEx dataset. We use *N* to indicate the number of individuals in each tissue that we have access to summary statistics from GTEx [19] and van de Bunt et al. (2015) study [25].

| chrom | pos. | rsID | Relevant Tissue | Target Gene |
| --- | --- | --- | --- | --- |
| 11 | 47293799 | rs10501320 | Pituitary ($N=87$)<br>Esophagus Muscularis ($N=218$)<br>Artery Tibial ($N=285$)<br>Esophagus Mucosa ($N=241$)<br>Islet ($N=118$)<br>Artery Tibial ($N=285$) | *ARHGAP1*<br>*C1QTNF4*<br>*MADD*<br>*MADD*<br>*MADD*<br>*MDK* |
| 11 | 72432985 | rs11603334 | Skin Sun Exposed Lower leg ($N=302$)<br>Pituitary ($N=87$)<br>Islet ($N=118$) | *ARAP1*<br>*PDE2A*<br>*STARD10* |
| 15 | 71109147 | rs1549318 | Adipose Visceral Omentum ($N=185$)<br>Cells Transformed Fibroblasts ($N=272$)<br>Ovary ($N=85$) | *LARP6*<br>*LARP6*<br>*LARP6* |

Many of the significant variants have CLPP values that are in the range where it is difficult conclude whether or not the causal variants are shared. However, we detect a large number of loci where the GWAS causal variants are clearly distinct from the causal variants in the eQTL data (Table 3). This includes several genes that can be excluded in all tissues tested (e.g., SEC22A at the rs11717195 FG-locus where there is non-colocalization).

**Table 3.** The loci where the causal variants between the eQTL and GWAS are different. We utilize FastingGlucose (FG) and FastingProinsulin (FP) phenotypes. Number of genes and tissues indicate the genes and tissues, respectively which we apply eCAVIAR for a GWAS risk variant. The complete list of genes and tissue are provided in Tables S2 and S3 for FG and FP phenotypes, respectively. eCAVIAR utilizes the marginal statistics of all variants in a locus obtained from GWAS and eQTL. We obtain each locus by considering 50 variants upstream and downstream of the reported variant. Then, we consider genes where at least one of the variants in the locus is significantly associated with the gene expression of that gene. Thus, for one GWAS variant, we can have multiple genes in one tissue that satisfy our condition.

| Phenotype | chrom | pos. | rsID | GWAS p-value | eQTL p-value<sup>1</sup> | # Gene | # Tissue |
| --- | --- | --- | --- | --- | --- | --- | --- |
| FG | 2 | 27741237 | rs780094 | 2.49E-12 | 2.95E-55 | 17 | 30 |
|  | 2 | 169763148 | rs560887 | 4.61E-75 | 1.36E-14 | 5 | 20 |
|  | 3 | 123065778 | rs11708067 | 8.72E-09 | 4.28E-42 | 5 | 34 |
|  | 9 | 4289050 | rs7034200 | 0.0001204 | 9.95E-12 | 8 | 7 |
|  | 10 | 113042093 | rs10885122 | 8.41E-11 | 7.73E-11 | 2 | 3 |
|  | 11 | 61571478 | rs174550 | 1.48E-08 | 1.03E-125 | 24 | 29 |
|  | 11 | 92708710 | rs10830963 | 1.26E-68 | 7.49E-06 | 7 | 6 |
| FP | 1 | 99177253 | rs9727115 | 5.285e-06 | 7.04e-16 | 3 | 12 |
|  | 10 | 114758349 | rs7903146 | 3.48e-18 | 7.92e-33 | 7 | 26 |
|  | 15 | 62383155 | rs4502156 | 3.80e-11 | 8.48e-14 | 7 | 15 |
|  | 17 | 2262703 | rs4790333 | 2.15e-08 | 5.39e-75 | 21 | 33 |

More interesting examples can be found amongst genes that colocalize in one tissue yet can be excluded in many others. An example of this can be found in *ADCY*5, also at the rs11717195 FG-locus. In pancreatic islet data, the GWAS variant itself colocalized with *ADCY*5 eQTLs, whereas eQTLs for the same gene show non-overlap with the GWAS association signal in several GTEx tissues. This suggests that the phenotype acts through a tissue-specific regulatory element active in islets yet inactive in other tissues.

For a majority of loci in which we identify a single variant causal for both GWAS and eQTL, our results implicate more than one target gene across the 45 tissues. eCAVIAR detects that three out of five colocalized variants in FG phenotype and all three variants in FP have multiple target genes. At rs7944584 (FG) / rs10501320 (FP) and rs1549318 (FP) there is support from other eQTL studies for causal roles for *MADD* in human pancreatic islets of Langerhans [25] and *LARP*6 in adipose tissue [22], respectively. Assessing the potential candidacy of these different implicated genes will require additional sources of information, such as Capture-C experiments [35], to demonstrate chromatin interactions between causal variant and gene promotor and/or in vitro function validation in relevant model systems. Even so, the current analysis leaves many loci where no colocalizing variant can be identified. The main reason for this is likely found in the limited power of eCAVIAR at the current sample sizes for the majority of tissues, especially for ones as pertinent to the phenotype as human islets (see Figure S10). Additional collection of samples will be necessary to overcome this hurdle and lead to further mechanistic insights.

## 3 Material and Methods

### 3.1 CAVIAR model for Fine-mapping

#### Standard GWAS and Indirect Association

We collect quantitative traits for *N* individuals and genotypes for all the individuals at *M* SNPs (variants). In this case, we collect data for one phenotype and gene expression of multiple genes. We assume that both the phenotype and the gene expression have at least one significant variant. To simplify the description of our method, we assume that the number of individuals and the pairwise Pearson’s correlations of genotype (LD) in both GWAS and eQTL are the same. In Appendix, we use a more general model where the number of individuals and LD in both GWAS and eQTL are not the same. Let *Y*^(*p*)^ indicate a (*N* × 1) vector of the phenotypic values where 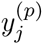 denotes the phenotypic value for the *j*-th individuals. We use *Y*^(*e*)^ to indicate a (*N* × 1) vector of gene expression collected for one gene of interest, for which there exists one significant variant associated with the gene expression of that gene. Let *G* indicate a (*N* × *M*) matrix of genotype information where *G_i_* is a (*N* × 1) vector of minor allele counts for all the *N* individuals at the *i*-th variant. In this setting *g_ji_* indicates the *j*-th element from vector *G_i_* that indicates the minor allele count for the *j*-th individual. In diploid genomes such as human, we can have three possible minor allele counts, *g_ji_* = {0,1,2}. We standardize both the phenotypes and the genotypes to mean zero and variance one, where the *X* is the standardized matrix of the *G*. Let *X_i_* denote a (*N* × 1) vector of standardized minor allele counts for the *i*-th variant. We assume “additive” Fisher’s polygenic model, which is widely used by GWAS community. In the Fisher’s polygenic model, the phenotypes follow a normal distribution. The additive assumption implies each variant contribute linearly to the phenotype. Thus, we consider the following linear model:

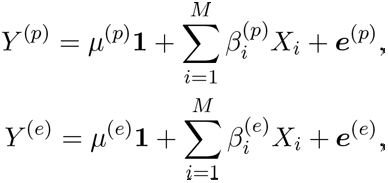

where *μ*^(*p*)^ and *μ*^(*e*)^ are the phenotypic and gene expression mean, respectively. Let 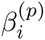 and 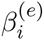 be the effect size of the *i*-th variant towards the phenotypes and gene expression, respectively. In addition, *e*^(*p*)^ and *e*^(*e*)^ are the environment and measurement error toward the collected phenotype and gene expression, respectively. In this model, we assume *e*^(*p*)^ is a vector of i.i.d. and normally distributed. Let 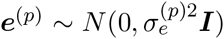 where 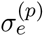 is a covariance scalar and ***I*** is a (*N* × *N*) identity matrix.

In our setting, we have the marginal statistics of *M* variants for phenotype of interest and the gene expression. Let 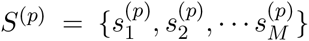 and 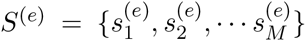 indicate the marginal statistics for the phenotype of interest and the gene expression, respectively. The joint distribution of the marginal statistics, given the true effect sizes, follows a multivariate normal (MVN) distribution and is similar to that found in previous work [18, 29–31]. Thus, we have:

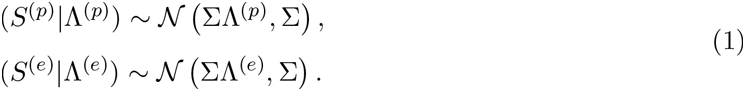

where Σ is the pairwise Pearson’s correlations of genotypes. Let 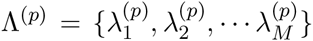 and 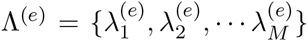 be the true standardized effect size for all the variants of desired phenotype and gene expression, respectively. The true effect size for a non-causal variant is zero and non-zero for causal variant. Let ΣΛ^(*e*)^ and ΣΛ^(*p*)^ be the LD-induced non-centrality-parameter (NCP) for desired phenotype and gene expression, respectively.

#### CAVIAR Generative Model for Single phenotype

We introduce a new variable *C*^(*p*)^ which is an (*M* × 1) binary vector. We refer to this binary vector as causal status. The causal status indicates which variants are causal and which are not. We set 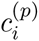 to be one if the *i*-th variant is causal and zero otherwise. In CAVIAR [18, 30], we introduce a prior on the vector of effect sizes utilizing the MVN distribution. This prior on the vector of effect sizes given the causal status vector is defined as follow:

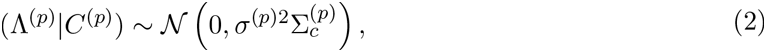

where 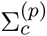 is a diagonal matrix and *σ*^(*p*)2^ is a constant which indicates the variance of our prior over the GWAS NCPs. We set *σ*^(*p*)2^ to 5.2 [18, 30]. The diagonal elements of 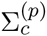 are set to one or zero where variants that are selected causal in *C*^(*p*)^ their corresponding diagonal elements in 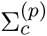 are set to one; otherwise, we set them to zero. Utilizing this prior as a conjugate prior, in CAVIAR, we compute the likelihood of each possible causal status. The joint distribution of the marginal statistics given the causal status is as follows:

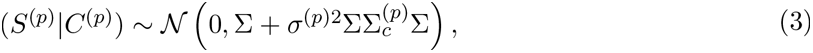

In a similar way, for the gene of interest that we perform eQTL, we have:

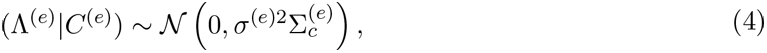

where 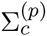 is a diagonal matrix and *σ*^(*e*)2^ is set to 5.2 [18, 30]. The diagonal elements of 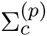 are set to one or zero where variants that are selected causal in *C*^(*e*)^ their corresponding diagonal elements in 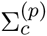 are set to one; otherwise, we set them to zero

### 3.2 eCAVIAR Computes the Colocalization Posterior Probability for GWAS and eQTL

Given the marginal statistics for GWAS and eQTL, which are denoted by *S*^(*p*)^ and *S*^(*e*)^, respectively, we want to compute the colocalization posterior probability (CLPP). CLPP is the probability that the same variant is causal in both studies. For simplicity, we compute CLPP for the *i*-th variant. We define CLPP for the *i*-th variant as 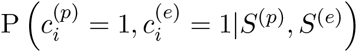, and we use *φ_i_* to indicate the CLPP for the *i*-th variant. We utilize the law of total probability to compute the summation probability of all causal status where the *i*-th variant is causal in both GWAS and eQTL and other variants can be causal or non-causal. Thus, the above equation can be extended as follows:

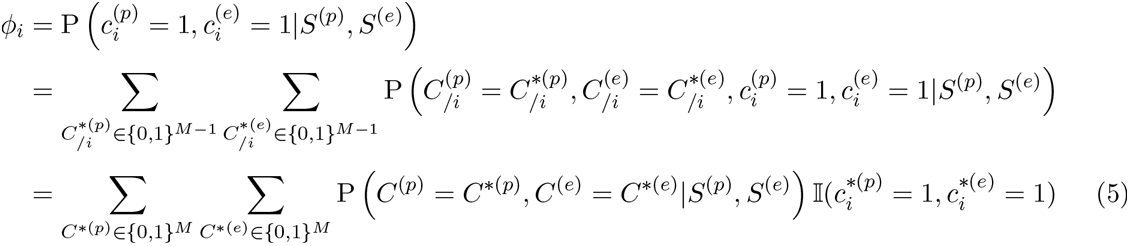

where 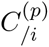 and 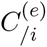 are causal status vectors for all the variants, excluding the *i*-th variant for the phenotype of interest and gene expression, respectively. Let 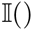 be an indicator function which is defined as follows:

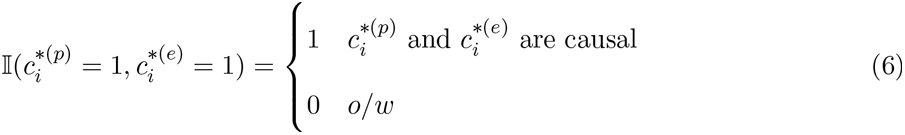

Utilizing the Bayes’ rule, we compute the CLPP as follows:

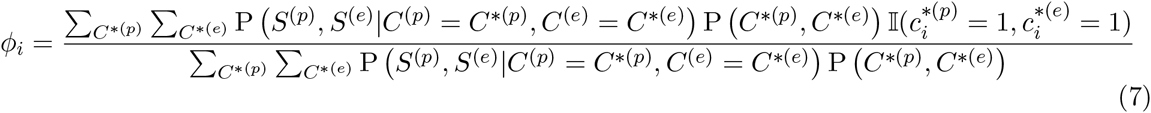

where P (*C*\*^(*p*)^,*C*\*^(*e*)^) is the prior probability of the causal status of *C*\*^(*p*)^ and *C*\*^(*e*)^ for the GWAS and eQTL respectively. We assume the prior probability over the causal status for the GWAS and eQTL are independent, P(*C*\*^(*p*)^,*C*\*^(*e*)^) = P (*C*\*^(*p*)^) P (*C*\*^(*e*)^). To compute the prior of causal status, we use the same assumptions that are widely used in the fine mapping methods [18, 30, 36], where the probability of causal status follows a Binomial distribution with the probability of variant being causal is equal to *γ*. Thus, this prior is equal to 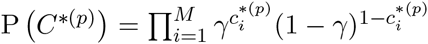 and *γ* is set to 0.01 [18, 37–39].

GWAS and eQTL studies are usually performed on independent sets of individuals. Furthermore, given the causal status of both the GWAS and eQTL, the marginal statistics for these two studies are independent. We have P(*S*^(*p*)^,*S*^(*e*)^|*C*\*^(*p*)^,*C*\*^(*e*)^) = P (*S*^(*p*)^|*C*\*^(*p*)^) P (*S*^(*e*)^|*C*\*^(*e*)^). Thus, we simplify the Equation (7), and the CLPP is computed as follows:

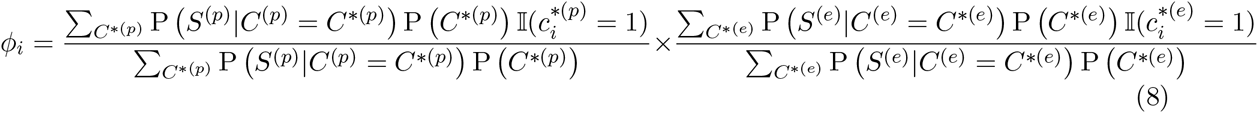

The above equation indicates the probability that the same variant is causal in both GWAS and eQTL is independent. This probability is equal to the multiplication of two probabilities: probability that the variant is causal in GWAS and the probability of the same variant is causal in the eQTL study. Thus, we compute the CLPP as, 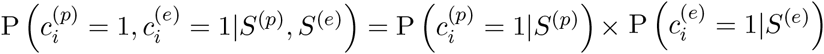 where 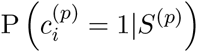 and 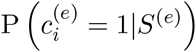 are computed from the first part and second part of Equation (8), respectively.

### 3.3 Detecting Target Genes and Relavent Tissues

In the previous sections, we describe the process of computing the CLPP score for each variant in a locus for a given eGene in a tissue. In this section, we describe a systematic way to detect the target genes and relavent tissues.

We compute the CLPP score for every GWAS significant variant. Thus, for a given GWAS variant, an eGene that has a CLPP score above the colocalization cut-off is considered a target gene. In addition, we consider tissues from which the target genes are obtained as the relevant tissues. Moreover, we can rank the relevant tissues and target genes for a given GWAS significant variant based on their CLPP scores. Thus, we utilize the magnitude of CLPP to rank the tissues and genes based on their importance for a given GWAS risk locus.

### 3.4 Generating simulated datasets

#### 3.4.1 Simulating Genotypes

We first simulated genotype data staring from the real genotypes obtained from European population in the 1000 Genomes data [26, 27]. In order to simulate the genotypes we utilize HAPGEN2 [33] software which is widely used to generate genotypes. We focus on the chromosome 1 and the GWAS variants that are obtained from the NHGRI catalog [40]. We consider 200-kb windows around the lead SNP to generate a locus. Then, we filter out monomorphic SNPs and SNPs with low minor alley frequency (MAF ≤ 0.01) inside a locus.

#### 3.4.2 Simulating Summary Statistics Directly from LD Structure

We generate LD matrix for a locus by computing the genotype Pearson’s correlations between each pair of variants. Then, we generate marginal summary statistics for each locus, assuming the marginal summary statistics follows MVN that is utilized in previous studies [18, 29–32]. We measure the strength of a causal variant based on NCP. We set the NCP of the causal variant in order to obtain a certain statistical power. The NCPs of the non-causal variants are set to zero. The statistical power is the probability of detecting a variant to be causal under the assumption that the causal variant is present. The statistical power is computed as follows:

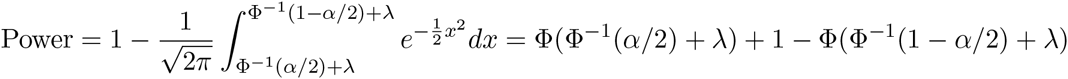

where *α* is the significant threshold. Moreover, Φ and Φ^−1^ denote the cumulative density function (CDF) and inverse of CDF for the standard normal distribution. In our experiment, the NCP is computed for the genome-wide significant level (*α* = 10^−8^). We use binary search to compute the value of NCP for a desired statistical power.

#### 3.4.3 Simulating Summary Statistics Utilizing Linear Additive Model

We utilize 100 variants in a locus to generate the simulated phenotypes from the simulated genotypes. We simulate the phenotypes assuming the linear additive model, which is as follows:

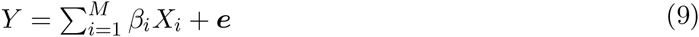

where 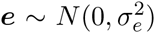. We generate the effect size of the causal variant from a normal distribution with mean zero and variance equal to 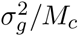 where *M_c_* indicates the number of causal variants in a locus. Furthermore, we set the effect size to zero for variants that are not causal. Thus, the effect size for each variant is simulated as follows:

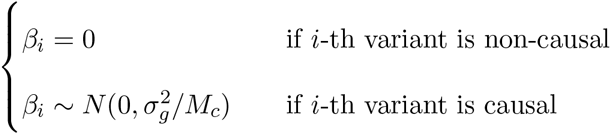

After simulating phenotype for all the individuals, we utilize linear regression to estimate the effect sizes and the marginal statistics for all the *M* variants in a locus. In our simulations, *M* is equal to 100.

## 4 Discussion

Integrating GWAS and eQTL provides insights into the underlying mechanism for genetic variants detected in GWAS. In this paper, we propose a quantity that can measure CLPP, the probability that the same variant is causal in both GWAS and eQTL studies, while accounting for the LD. Utilizing CLPP, we can identify target genes and relevant tissues. It is worth mentioning that we can use epigenomic data (e.g., NIH Roadmap Epigenomics [41]) to detect relevant tissues as an orthogonal analysis compared to using eCAVIAR. Moreover, eCAVIAR can detect loci where the causal variants are different between the two studies with high confidence. We observe from our analysis that, in most cases, GWAS risk loci and eQTL are different.

As most GWAS loci were discovered to lie outside of coding regions, it is implicitly assumed that these implicated loci will affect the regulation of genes. However, our results produce a lower than expected number of variants colocalized between both GWAS and eQTL studies. This points to a more complicated relationship between gene regulation and disease. It is likely that future studies will shed some light to explain this observation.

One conjecture is that the GWAS loci in fact do affect expression, but are secondary signals compared to the stronger associations found in current eQTL studies. As eQTL studies include an increasing number of individuals, we will be able to prove or disprove this conjecture. Furthermore, the heterogeneity of tissues may render it hard to detect eQTLs specific to a disease-relevant cell type that comprises only a fraction of the tissue. A second possibility is that GWAS variants affect other aspects of gene regulation such as splicing, or regulation at a level other than transcription regulation. Several studies have shown that alternative splicing may explain the causal mechanism of complex disease associations (e.g., a variant associated with multiple sclerosis that leads to exon skipping in *SP*140 [42]). Methods that identify variants associated with differences in relative expression of alternative transcript isoforms or exon junction abundances are being applied to the latest version of GTEx data [43, 44]. As we obtain more functional genomics information and are able to measure quantities such as protein abundance, we will be able to systematically catalogue variants that affect regulation at levels other than transcription. A third possibility is that GWAS loci are eQTL loci only in certain conditions, such as development, where expression levels are not typically measured. Regardless, our study demonstrates strong evidence in support of the idea that most GWAS loci are not strong eQTL loci and that the mechanism of GWAS loci affecting gene regulation is more complicated than we expected.

Broadly, we identify an analogy between colocalization and fine-mapping methods. Fine-mapping methods can be categorized into three main classes. One class relies on just the computed marginal statistics that are obtained from GWAS or eQTL. In this class of methods, the probability that a variant is causal depends on the rank of a variant, which is obtained from the marginal statistics. Recently, Maller et. al [45] have proposed a new fine-mapping method that utilizes the Bayes factor. This method provides results similar to those produced by approaches that rank variants based solely on their marginal statistics. Maller et. al [45] method for fine-mapping is similar in nature to COLOC [14], which is used for colocalization. The second class of methods is based on a conditional model where we re-compute the marginal statistics of all variants by conditioning on variants selected as causal. The conditional method for fine-mapping and RTC [13] have some similarity in nature. The third class of methods is CAVIAR [18, 30], CAVIARBF [46], and FINEMAP [36], which assumes a presence of more than one causal variant in a region. These probabilistic based methods use the MVN distribution. In these methods, we detect a set of variants that can capture all the causal variants with a predefined probability. eCAVIAR is analogous in process to CAVIAR, CAVIARBF, and FINEMAP. However, eCAVIAR and CAVIAR-like methods try to solve different problems. CAVIAR-like methods (CAVIARBF and FINEMAP) are designed to perform fine-mapping. CAVIARBF is based on the CAVIAR statistical model that utilizes Bayes factor to detect the causal set. FINEMAP is based on the CAVIAR statistical model that utilizes sampling techniques to speed up the computational process of detecting the causal set.

eCAVIAR is a probabilistic method that integrates GWAS and eQTL signals to detect biological mechanisms. eCAVIAR has several advantages over prior approaches. First, it can account for multiple causal variants in any given locus. Second, it leverages summary statistics without accessing the raw individual data. In addition, eCAVIAR can provide confidence levels for the colocalization of a GWAS risk variant. Utilizing the confidence level, we can categorize a variant to three categories: variants which colocalize, variants which do not colocalize, and variants which are ambiguous to detect their colocalization status for the current data. High-throughput technologies have made it possible to obtain multi-tissue eQTL studies. Leveraging multi-tissue eQTL studies such as GTEx and eCAVIAR can advance discovery of new biological mechanisms for GWAS risk loci. eCAVIAR can be extended to utilize functional annotations to improve our results. This can be done similar to PAINTOR [34] and RiVIERA-beta [47] that incorporate functional annotations to improve fine-mapping results.

## Acknowledgments

FH, JWJJ, MB and EE are supported by National Science Foundation grants 0513612, 0731455, 0729049, 0916676, 1065276, 1302448, 1320589 and 1331176, and National Institutes of Health grants K25-HL080079, U01-DA024417, P01-HL30568, P01-HL28481, R01-GM083198, R01-ES021801, R01-MH101782 and R01-ES022282. EE is supported in part by the NIH BD2K award, U54EB020403. MvdB is supported by a Novo Nordisk postdoctoral fellowship run in partnership with the University of Oxford. AVS and XL are supported by contract HHSN268201000029C (Broad Institute). SS was supported in part by NIH grant R00-GM 111744–03. We acknowledge the support of the NINDS Informatics Center for Neurogenetics and Neurogenomics (P30 NS062691).

## 6 Web Resources

eCAVIAR is available http://genetics.cs.ucla.edu/caviar/

## Appendix

## CAVIAR and eCAVIAR General Models where GWAS and eQTL Have Different Number of Individuals

### Standard Association Test in GWAS

We assume that there are *N*^(*p*)^ individuals in the GWAS for the phenotype of interest where we collect the phenotypic values of a quantitative trait for all the individuals. Moreover, we collect the genotypes of all the individuals for *M* variants. Let *Y*^(*p*)^ be a (*N*^(*p*)^ × 1) vector of phenotypic values, which is obtained from GWAS. Let *G*^(*p*)^ be a (*N*^(*p*)^ × *M*) matrix of genotypes where 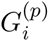 is (*N*^(*p*)^ × 1) vector of the minor allele count for the *i*-th variant. We use 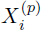 to indicate the standardized vector of the minor allele count for the *i*-th variant, 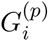, where 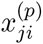 is the standardized genotype of the *i*-th variant for the *j*-th individual. We assume both the phenotypes and genotypes are standardized. We standardize a vector to make the mean and the variance to be equal to zero and one, respectively. Thus, we have 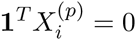 and 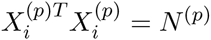 that we can show as follows:

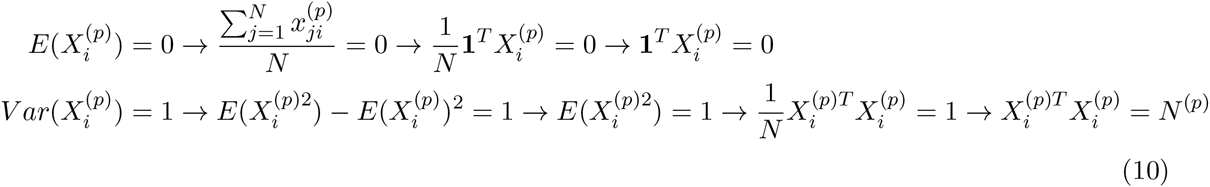

We assume the “additive” Fisher’s polygenic model. In this model, each variant has small effect towards the phenotype where these effects are linear and additive. Thus, we have:

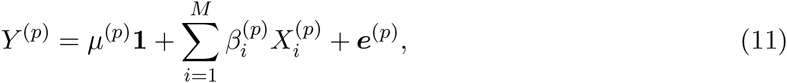

where *μ*^(*p*)^ is the population mean for the phenotype, 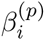 is the effect of the *i*-th variant towards the phenotype, and *e*^(*p*)^ is the environment and measurement noise that we assume follows a normal distribution, 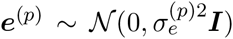 where 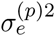 is covariance scalar. As mentioned in the method section, we test the significance of each variant one at a time. Moreover, we assume the c-th variant is causal. Thus, the model we have is as follows:

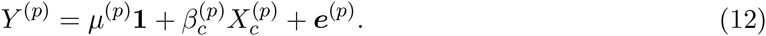

To ease our notations, we utilize the fact that both phenotype and genotypes are standardized. Thus, the phenotypes follow a normal distribution with mean equal to 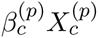 and variance of 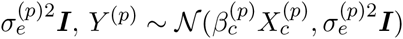. To estimate the effect size, we utilize the maximum likelihood. The likelihood is computed as follows:

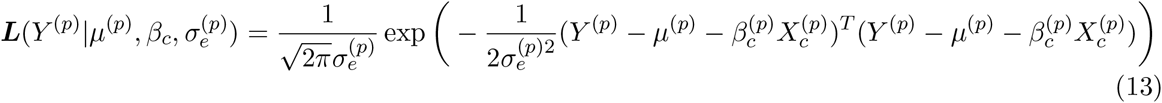

we compute the optimal effect size that maximizes the above likelihood, by computing the derivate of likelihood and setting it to zero. As a result the optimal effect size is computed as follows:

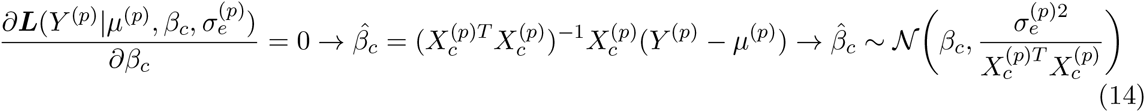

We compute the marginal statistics by dividing the estimated effect size by the standard deviation of the estimated effect size. Thus, we have:

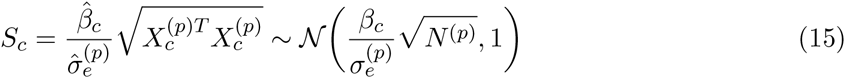

where 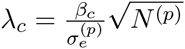 is the NCP of *c*-th variant, which is the causal variant.

### Indirect Association Test in GWAS

We assume there are two variants where c-th variant is causal and *i*-th variant is not causal. To estimate the effect size we use the same testing model that we show in the previous section. Thus, we have the following model:

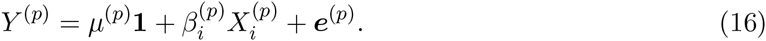

where we maximize the likelihood function to obtain the optimal effect sizes. The optimal effect size for the *i*-th variant is as follows:

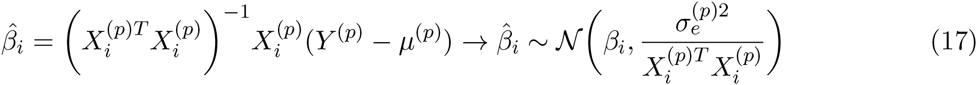

We compute the marginal statistics similar to the previous section as follows:

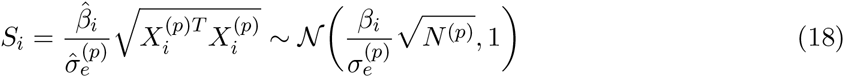

The variance of the marginal statistics is one. Thus, correlation and covariance of marginal statistics are equal. We compute the covariance of the marginal statistics between the causal variant and non-causal variant. We compute this correlation as follows:

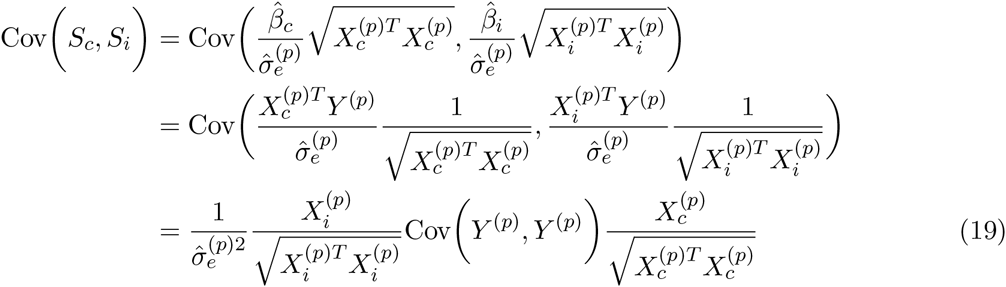

Using the Slutsky’s theorem and the fact that the number of individuals in a study is large enough, 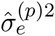 approaches Var(*Y*^(*p*)^). Thus, we have:

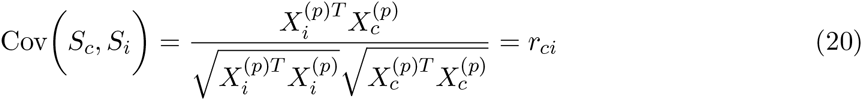

This indicates that the correlation between the marginal statistics of two variants is equal to their genotype correlation. This result is known from our previous studies [18, 30, 48].

### Standard Association Test in eQTL

We assume in the eQTL study, we collect the gene expression of multiple genes for *N*^(*e*)^ individuals. Let superscript (*p*) and (*e*) indicate variable related to GWAS and eQTL, respectively. Let *Y*^(*e*)^ indicate the gene expression level of all the *N*^(*e*)^ individuals for one gene. We consider one gene to ease the presentation of the method. We use the same model, which is the “additive” Fisher’s polygenic model, that we illustrate in the previous section for GWAS.

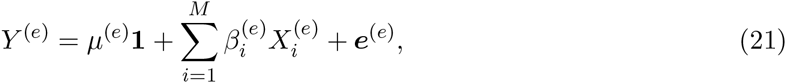

where *μ*^(*e*)^ is the population mean for the gene expression for the gene of interest, 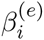 is the effect of the *i*-th variant towards the gene expression, and *e*^(*e*)^ is the environment and measurement noise which follow a normal distribution, 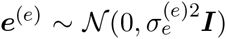 where 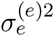 is covariance scalar. As mentioned in the method section, we test the significant of each variant one at a time. Similarly, we assume the c-th variant is causal. Thus, the model we have is as follows:

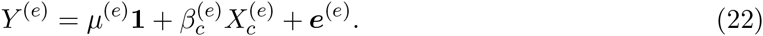

The optimal estimated effect size for eQTL is similar to optimal estimated effect size for GWAS.

### CAVIAR model for GWAS and eQTL

We know that the covariance between the estimated effect size of two variants is equal to their genotype correlation. Furthermore, the mean of the marginal statistics of the non-causal variants is equal to the mean of marginal statistics of the causal variants scaled by the genotype correlation. Thus, we have:

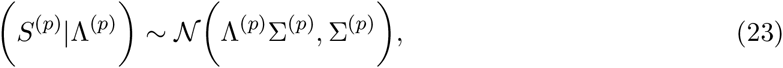

where Σ^(*p*)^ matrix is the pairwise genotype correlations obtained from GWAS. For the eQTL study, we obtain similar equation for the joint marginal statistics that is as follows:

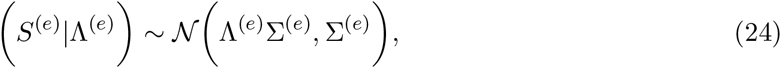

where Σ^(*p*)^ matrix is the pairwise genotype correlations obtained from the eQTL study. We consider Λ^(*p*)^ and Λ^(*e*)^ are the true effect size vectors for GWAS and eQTL studies respectively. True effect sizes are non-zero for causal variants and are zero for the non-causal variants. Moreover, we consider Λ^(*p*)^Σ^(*p*)^ and Λ^(*e*)^Σ^(*e*)^ are the LD-induced effect sizes for GWAS and eQTL studies, respectively.

We introduce a MVN prior over the true effect size vectors. The true effect sizes for variants are independent and for causal variants are non-zero. Thus, we have the following prior:

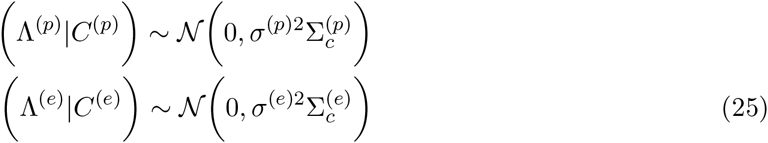

where 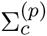 is a diagonal matrix and *σ*^(*p*)2^ is set to 5.2 [18, 30], which indicates the variance of our prior over the GWAS effect sizes. The diagonal elements are set to one or zero where variants which are selected causal we set the corresponding diagonal elements to one; otherwise, we set them to zero.

Utilizing the conjugate prior, we can combine Equations (23) and (25) to obtain the joint distribution of the marginal statistics given the causal status vector. This distribution is as follows:

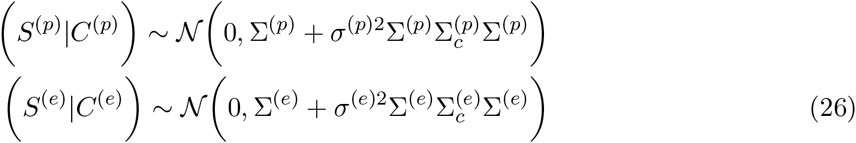

To show the correctness of the above equations, we utilize the law of total expectation and law of total variance. Given two random variables A and B, the law of total expectation is as follows:

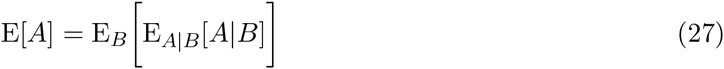

Let *A* = (*S*^(*p*)^|*C*^(*p*)^) and *B* = Λ^(*p*)^, we can compute the mean of the marginal statistics given the causal status as follow:

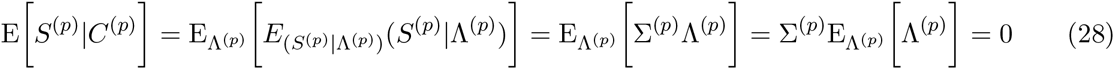

To compute the variance of the joint distribution of the marginal statistics given the causal status, we use the law of total variance that is as follows:

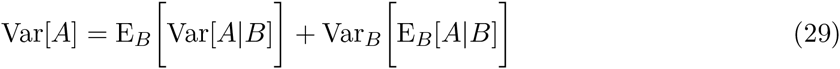

Thus, we compute the variance of joint distribution as follows:

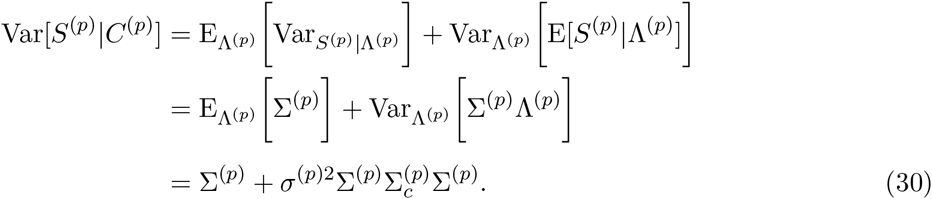

## Footnotes

1 The *p*-value of the most significant variant in eQTL among all genes and all tissues

